# Plasticity of the xylem vulnerability to embolism in poplar relies on quantitative pit properties rather than on pit structure

**DOI:** 10.1101/2020.05.14.096222

**Authors:** Cédric Lemaire, Yann Quilichini, Nicole Brunel-Michac, Jérémie Santini, Liliane Berti, Julien Cartailler, Pierre Conchon, Éric Badel, Stéphane Herbette

## Abstract

Knowledge on variations of drought resistance traits are needed to predict the potential of trees to acclimate to coming severe drought events. Xylem vulnerability to embolism is a key parameter related to such droughts, and its phenotypic variability relies mainly on environmental plasticity. We investigated the structural determinants controlling the plasticity of vulnerability to embolism, focusing on the key elements involved in the air bubble entry in a vessel, especially the inter-vessel pits. Poplar saplings (*Populus tremula x alba*) grown in contrasted water availability or light exposure exhibited differences in vulnerability to embolism in a range of 0.76 MPa. We then characterized the structural changes related to qualitative and quantitative pit characteristics, from the pit structure to the organization of xylem vessels, using different microscopy techniques (TEM, SEM, light). X-ray microtomography analysis allowed observing the vessel vulnerability and testing some of the relationships between structural traits and vulnerability to embolism inside the xylem. The pit ultrastructure did not change, whereas the vessel dimensions increased with vulnerability to embolism and the grouping index and fraction of inter-vessel cell wall decreased with vulnerability to embolism. These findings holds when comparing trees or when comparing vessels inside the xylem. These results evidenced that plasticity of vulnerability to embolism occurs through changes in the quantitative pit properties such as pit area and vessel grouping rather than on the pit structure.

## Introduction

According to the cohesion-tension theory (Steudle 2001), the water columns in the xylem are under tension, a metastable state. When tension forces increase during droughts, the water columns are more prone to break, because of cavitation: vapour bubbles invade the impacted vessels and hence make them embolized as they are not functional anymore, leading to a loss of xylem conductance. When the loss of conductance reaches a high threshold limit, the above organs are not supplied with water anymore leading to death (Barigah et al. 2013). For woody species, drought-induced death is more likely due to xylem hydraulic failure (Anderegg et al. 2015, 2016) caused by embolism in the xylem conduits, even if over process can also contribute to this death (Hammond et al. 2019) such as the carbon starvation (Hartmann et al. 2015).

A global analysis pointed out the narrow hydraulic safety margin at which woody species usually operate (Choat et al. 2012); inferring that research is needed on the variability for vulnerability to embolism. Within-species variability for vulnerability to embolism has been shown for many tree species (e.g. Martínez-Vilalta et al. 2009; Herbette et al. 2010). The genetic variability for this trait is rather limited in both natural populations (Lamy et al. 2011, Wortemann et al. 2011) and cultivated species (Jinagool et al. 2015, 2018). This trait would be genetically canalized (Lamy et al. 2012) such that it varies mainly via plasticity due to environmental factors (Herbette et al. 2010). Plasticity of vulnerability to embolism was reported mainly under water stress, with wood formed under drier conditions that tends to be less vulnerable (Awad et al. 2010, Fichot et al. 2010, Plavcová and Hacke 2012). Other conditions such as shade or fertilization were associated to an increase in vulnerability to embolism (Cooke et al. 2005; Barigah et al. 2006; Plavcová and Hacke 2012). However, information is scarce on the determinants that control the plasticity of vulnerability to embolism. The structural determinants need to be deciphered first, before searching for their genetic control, as it can be complex to decipher the role of candidate genes (Allario et al. 2018).

In angiosperms, water flows in the xylem vessels through bordered pits. These pits are cavities in the secondary cell wall that allow the inter-vessel water flow while they prevent air seeding from air-filled vessels to water-filled ones. Inter-vessel pits have been identified as the key structures for vulnerability to embolism (Lens et al. 2013). Thus, we assume that the acclimation of vulnerability to embolism to environmental conditions implies changes in the pit properties. The quantitative and qualitative pit properties that relate to vulnerability to embolism have to be investigated at the xylem, vessel and pit levels (Lens et al. 2013). The key role of the pit ultrastructure in vulnerability to embolism has been evidenced in several studies (e.g. Lens et al. 2011, Tixier et al. 2014), especially the pit membrane thickness (Jansen et al. 2009). The vulnerability to embolism is also depending on the different quantitative pit parameters such as the pit area or the vessel connectivity (e.g. Lens et al. 2011). Zimmermann and Jeje (1981) pointed out that the hydraulic vulnerability could be related to the vessel volume that varies depending on both their diameter (Tyree et al. 1994) and their length (Scholz et al. 2013a). The relationship between vulnerability to embolism and pit properties has been intensively studied at the inter-specific level, whereas the determinants of the plasticity of vulnerability to embolism remains poorly investigated or unclear at the intraspecific level. For example in poplar, shading caused an increase in vulnerability to embolism with a decrease in both pit membrane thickness and vessel diameter (Plavcová et al. 2011) whereas a reduced watering caused a decrease in vulnerability to embolism linked with a decrease in vessel diameter (Awad et al. 2010).

In this work, we investigated the relationship between the plasticity of vulnerability to embolism and changes in the structure related to pit properties at different anatomical levels on young poplars (*Populus tremula x alba*). We grew sapling poplar clones under three contrasted environmental conditions for two factors (water availability and light exposure) known to induce vulnerability to embolism plasticity. Then, their xylem anatomy was analysed in relation to the changes in vulnerability to embolism using different approaches. Transmission Electron Microscopy (TEM) allowed investigations on the bordered pit ultrastructure. Parameters related to the pit-field were measured using Scanning Electron Microscopy (SEM). We also measured quantitative pit parameters related to vessel dimensions and vessel connectivity using light microscopy and silicon injections. Then, a local approach using direct observation of embolism spreading at the vessel level by X-ray microtomography allowed us to analyse some relationships between the hydraulic network structure and the vulnerability to embolism inside the xylem.

## Materials and Methods

### Plant material and growth conditions

#### Plant Material

Saplings of hybrid poplar (*Populus tremula x alba* clone INRA 717-1B4) were multiplied clonally in vitro on Murashige and Skoog medium on December 2016. Plantlets were grown in hydroponic solution on February 2017 in a controlled environment room: 16 h daylight at 21-22 °C, 40 µmol.m^−2^.s^−1^ and 18-19 °C night, at 70 ± 10 % relative humidity. On March 2017, plants were transferred in 1 Litre pots filled with potting soil (Humustar Terreaux, Champeix, France) with a composition of 25 % brown peat, 40 % blond peat and 35 % pine barkdust. The pots were placed in a greenhouse at the INRA research station of Clermont-Ferrand, France (site of Crouël; N 45°77′, E 3°14′; 300 m *a.s.l.*). After 20 days, plants were transferred in 10 L pots filled with potting soil. Pots were regularly watered at soil field capacity. Each pot weighted 6.4 ± 0.4 kg. Ten days later, the specific experimental growth conditions were applied (see next). After one month of growth, stems were cut at 50 cm height. The growth of a new apical bud occurred in May 2017, and any additional bud was removed. Thus, a single stem completely grew under the new environmental conditions.

#### Experimental setup

Plants were split in three groups submitted to different growth conditions: (i) “Control” plants grew under full sunlight and watered at soil field capacity; (ii) “Droughted” plants grew under full sunlight and watered at only 25-30 % of soil field capacity; (iii) “Shaded” plants shaded by a shadehouse that intercepts 30 % of incident light and watered at soil field capacity. For the nine Droughted plants, an irrigation at 25-30 % of soil field capacity was kept constant in each pot individually using balances and valves for irrigation as described in Niez et al. (2019). We measured the light interception by the shadehouse by comparing for two months the light intensities recorded between two sensors (PAR/CBE 80, Solems, Palaiseau, France) placed inside the shadehouse and two sensors placed outside. The level of water stress was set to be the most restrictive while allowing growth to produce acclimatized xylem and enough material plant for further analyses. The stem diameter was continuously measured using a LVDT sensor (Linear Variable Differential Transformer) on 3 Droughted, 2 Control, 3 Shaded plants. Plant height was also measured regularly using a tape measure.

The day before sampling, predawn water potential (*Ψ*_pd_) were measured on all plants 1 hour before the sunrise using a pressure chamber (1505D, PMS Instrument, Albany, OR, USA, Scholender et al. 1965). The midday water potential (*Ψ*_mid_) was measured at the solar noon, between 12h00 and 14h00 the same day.

#### Sampling protocol

The sampling was performed on 28 August 2017. Plants were cut at 20 cm height. The plant shoot was immerged underwater and the 30 cm of the top were removed as they have few developed xylem. Then, the following stem segments were sampled, from basal to apical direction:

i. the 30 cm long basal part of the stem was removed because it was not fully grown under acclimating conditions;
ii. the first 50 cm long of the newly developed stem under the acclimation conditions was wrapped in wet paper in a plastic bag and stored at 4 °C before measurements of vulnerability to embolism and vessel length;
iii. the above segment of 6 cm long was devoted to microscopy analyses. It was split into three subsamples using a razor blade: two segments of 1 cm long were prepared for light microscopy and TEM, and a third segment of 4 cm long was prepared for SEM;
iv. when the stem was long enough, an additional segment of 50 cm long was wrapped in wet paper in a plastic bag, and stored at 4 °C for measurements of specific conductivity (*K*_S_) and for additional measurements of vulnerability to embolism;
v. the last 10 cm long was kept wrapped in humid paper for a native embolism measurement performed during the sampling day.

Leaves were sampled under water and the mean leaf area (LA) per plant was measured in the day using an area-meter (Li-3100c, Li-Cor Biosciences, Lincoln, NE, USA).

After the sampling, plants were kept in the greenhouse, during the winter 2017. On March 2018, they started growing, still under the same environmental conditions as described above, and on July 2018 we performed a new sample collection: plants were cut at 25 cm height. Then the 30 cm long basal part of the stem were cut underwater. A 50 cm long sample was wrapped in wet paper and stored in a plastic bag at 4 °C for measurements of specific conductivity (*K*_S_).

### Hydraulic traits

#### Vulnerability to embolism

The 50 cm long stem segment was cut underwater at 43 cm long using a razor blade. Then, the vulnerability to embolism was assessed using the Cavitron technique (Cochard 2002, Cochard et al. 2005). The centrifugal forces increase water tension in branch segment and allows at the same time measurement of the loss of conductance using a reference ionic solution of 10 mM KCl and 1 mM CaCl_2_ (Cochard et al. 2009). A vulnerability curve was built by plotting the percentage loss xylem conductance (PLC) *vs.* xylem water pressure (*P*). A sigmoidal function was used to fit each curve using the equation 1 (Pammenter and Willigen 1998).

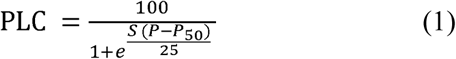

Where *P*_50_ is the pressure causing 50 % loss of conductance, and *S* the slope of the curve at this point.

#### Specific conductivity

Stem segments of 50 cm long were cut underwater at a length (*L*_stem_) of 40 cm long using a razor blade for Droughted (n = 8), Control (n = 9) and Shaded (n = 9) plants. The apical end of the sample was sealed to a tubing system (polytetrafluoroethylene film) and plugged to an embolism meter (Xyl’em, Bronkhorst, Montigny les Cormeilles, France). The initial conductance (*K*_i_) is then measured under low pressure (2 to 7 kPa) using a solution of 10 mM KCl and 1 mM CaCl_2_. The xylem area *A*_X_ of the distal end of the sample was measured on a cross section using a scanner (V800, Epson, Nagano, Japan). The measurement of *A*_x_ was performed on the scanned image using the ImageJ software (version v.1.52c) (Schneider et al. 2012). The Specific Conductivity *K*_S_ was defined according to equation 2.

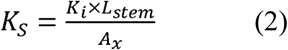

#### Native Embolism

The native embolism of the stem segments of 10 cm long were measured the day of their harvest for Droughted (n = 9), Control (n = 5) and Shaded (n = 6). Each sample was cut underwater using a razor blade to a length of 8 cm. Then, the initial conductance (*K*_i_) was measured under low pressure (2 to 7 kPa) with the same method and the same solution as specific conductivity. Then, the sample was flushed with the same solution two times for 5 min under high pressure (0.1 to 0.2 MPa) in order to remove embolism. A new measurement of conductance without embolism gives the maximum conductance (*K*_max_) of the sample. The native embolism was calculated according to the equation 3.

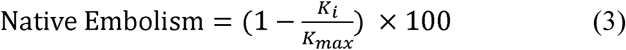

### Light microscopy

Samples of 1 cm long were cut into 3 × 3 mm^2^ blocks then they were immersed in Karnvosky’s fixative solution under vacuum for 30 min, then stored at 4 °C in the fixative solution up to the next step. Then, they were dehydrated in an ethanol series (50, 70, 80, and 95 %) and embedded in LR White medium. Transverse slices of 2 to 3 µm thick were cut using an ultramicrotome (Om U2, Reichert, Vienna, Austria). Sections were stained with 1 % (w/v) toluidine blue, washed 4 times with water and mounted in Eukitt (Sigma-Alrich, St-Louis, MO, USA). Images were processed using a microscope (Zeiss Axio Observer Z1), a digital camera (AxioCam MRc) and Zen imaging software system (Zeiss, Jena, Germany). Image analyses were performed using ImageJ software. The vessel diameter (*D*_v_) was estimated to be the diameter of the circle having the same area as the vessel lumen (for the symbols, see Table 1). The contact fraction (*F*_c_) was measured for each vessel as the ratio of wall length shared with other vessels over the vessel perimeter. The grouping index (GI) was measured as the mean number of vessels per group and the solitary index (SI) as the ratio of the number of solitary vessels to the total number of vessels. These parameters were measured for each individual slice containing a mean of 850 vessels, for Droughted (n = 9), Control (n = 5) and Shaded (n = 6) plants.

**Table 1:**
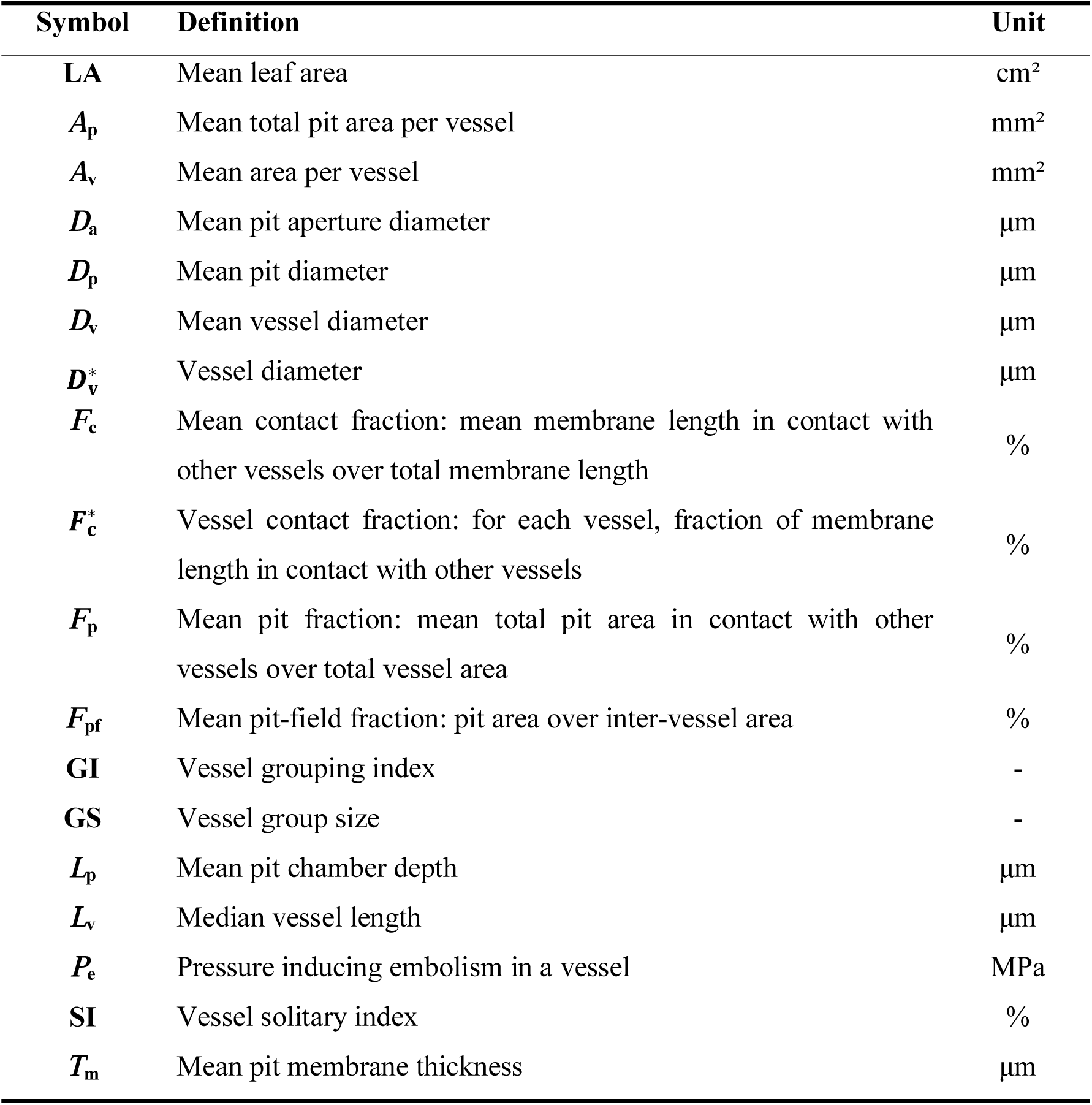
Meanings of the symbols.

### Vessel length

The vessel length was measured by the silicon injection method (Sperry et al. 2005, Scholz et al. 2013b) on the samples used for Cavitron technique, after five months of drying at room temperature. A fluorescent optical brightener (CAS number: 7128-64-5, Sigma-Aldrich, St-Louis, MO, USA) was mixed in chloroform (1 % w/w) and added to a volume of silicon (BLUESIL RTV-141 A, Bluestar Silicones, Lyon, France) with a proportion of one drop of solution per gram of silicon. A Silicone hardener (BLUESIL RTV-141 B, Bluestar Silicones) was added to the mixture in 1:10 proportion. The mixture was then injected under pressure (30 to 40 kPa) basipetally in the stem sample using a pressure chamber during at least 8 hours. After silicone hardening (3 days at room temperature), the samples were cut 5 mm far from the injection point; then every 20 mm. For each segment, one 25 µm thick slice was cut using a rotary microtome (RM2165, Leica Microsystems, Wetzlar, Germany). Cross section slices were dyed with Astra Blue and mounted with a Lugol’s iodine solution.

Images were obtained using a fluorescence microscope (Axio Observer Z1) equipped with a 300 to 400 nm band pass excitation filter, a digital camera (AxioCam 506), Zen imaging software system (Zeiss, Jena, Germany) and analysed using the ImageJ software. Fluorescent vessels highlighted the open vessels, while white light allowed counting the total number of vessels. The decrease of the ratio of open vessels (*N*_x_) (*i.e.* fluorescent vessels) to the total number of vessels (*N*_0_) over the distance (*x*) from the end of the sample followed a Weibull function (equation 4) where *k* is the best-fit extinction coefficient (Cohen et al. 2003).

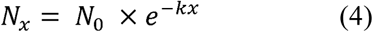

The fraction of conduits of length *x* (*P*(*x*)) is obtained by multiplying *x*/*N*_0_ to the second derivative of equation 4 (Wheeler et al. 2005):

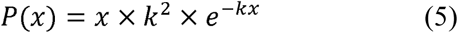

The continuous cumulative function of vessel length (*L*v) probability is a function given in the equation 6.

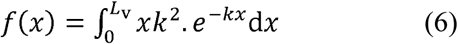

When this cumulative function is equal to 0.5, this gives the median value of vessel length (*L*_v_) (equation 7).

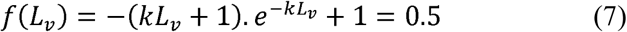

The solution of the equation 7 gives the median vessel length *L*_*v*_ = 1.678/*k*. This vessel length was estimated for 7 Droughted, 5 Control and 5 Shaded stem samples.

### Transmission Electron Microscopy

Fresh samples of 1 cm long were cut into 2 to 4 mm^3^ blocks, immersed in Karnvosky’s fixative solution under vacuum for 30 min, then stored at 4 °C in the fixative solution for 3 weeks. Blocks were recut into 1 to 2 mm^3^ pieces, then they were secondary fixed for 4 hours at ambient temperature in a 0.1M phosphate-buffered osmium tetroxide solution (1 %), pH 7.4. Then, they were dehydrated in an ethanol series (25, 50, 70, 100, and 100 %) and embedded in Epoxy resin using Epoxy medium kit (Sigma-Aldrich, St-Louis, MO, USA). Then, ultra-thin sections (60-90 nm) were cut using an ultramicrotome (PowerTome PC, RMC Boeckeler, Tucson, AZ, USA). The sections were placed on 200- and 300-mesh copper grids and stained with contrast solutions: UranyLess (Delta Microscopies, Mauressac, France) and lead citrate. Sections were observed using a transmission electron microscope (H-7650, Hitachi High-Technologies Corporation, Tokyo, Japan) at a voltage of 80 kV. Measurements of pit features were performed on images with pits showing two apertures. Pits were characterized for their diameter (*D*_p_), their aperture diameter (*D*_a_), their depth of pit chamber (*L*_p_) and their membrane thickness (*T*_m_). For each pit, *D*_a_ was the mean of two measurements while *L*_p_ and *T*_m_ were the mean of four measurements. Pit features were measured for five individual trees for each growth condition, with at least 10 pits measured per individual tree.

### Scanning Electron Microscopy

Fresh samples were fixed in glutaraldehyde 3 % fixative solution and stored at 4 °C for at least 1 month. Samples of 4 cm long were cut in a longitudinal way and were dehydrated in an ethanol series (30, 50, 75, and 100 %). After dehydration, samples were immerged in a 1:1 solution hexamethyldisilazane (HMDS) + ethanol 100 % for 30 min and immerged in pure HDMS for 30 min. After air drying overnight under a hood, the samples were mounted on aluminium stubs with carbon double-sided adhesive disks, coated with gold/palladium in a sputter coater (SC7640, Quorum Technologies Ltd, Newhaven, U.K.), and finally observed using a scanning electron microscope (S-3400N, Hitachi High-Technologies Corporation, Tokyo, Japan) at a voltage of 5 kV. The portion of area covered by bordered pits in each inter-vessel pit-field (*F*_pf_) was measured using the ImageJ software. Five samples were measured per growth condition, and seven pit-fields were characterized per sample.

### Calculation of supplemental hydraulic and structural traits

Theoretical conductivities (*K*_h_) of all samples characterized for light microscopy were calculated according to Scholz et al. (2013b) and converted into mol.s^−1^.MPa^−1^.m^−1^ (equation 8).

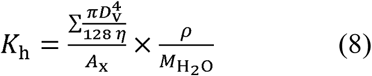

with η the viscosity index of water (1.002 × 10^−9^ m^4^.MPa^−1^.s^−1^ at 20 °C), ρ the density of water (9.982 × 10^5^ g.m^−3^), *M*_H2O_ the water molar mass (18.0 g.mol^−1^) and *A*_x_ the xylem area.

The pit fraction (*F*_p_) was defined as the product of the pit-field fraction (*F*_pf_) and contact fraction (*F*_c_) (equation 9).

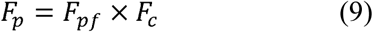

The pit fraction was measured on five individual trees for each growth condition.

The vessel area (*A*_v_) was calculated as the area of a cylinder according to the equation 10.

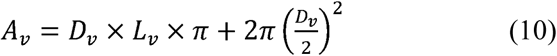

It was measured for 7 Droughted, 5 Control and 5 Shaded individuals.

The pit area per vessel (*A*_p_) was calculated as the product of the two above-cited traits (equation 11).

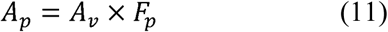

It was measured for 4 Droughted, 5 Control and 4 Shaded individuals.

Xylem water potentials at the onset of xylem embolism (*P*_12_) and at full embolism (*P*_88_) were calculated using equation 12 and 13 respectively (Domec and Gartner 2001), with *P*_50_ and *S* from equation 1.

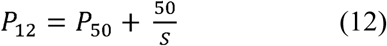

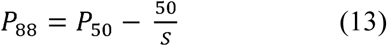

### Measurement of individual vessel vulnerability to embolism using X-Ray microtomography

Four stem segments from Droughted and Control plants were sampled and prepared in the same condition as for vulnerability to embolism measurements. We used the techniques described in Cochard et al. (2015). Segments were cut underwater at 43 cm long using a razor blade, sealed in liquid paraffin wax in order to prevent dehydration during the microtomography scans. A first 21 min scan was acquired using a X-ray microtomography system (Phoenix nanotom, General Electric, Boston, MA, USA) at the centre of the segment as described below to reveal the native state of embolism in each shoot. The field-of view was 7.8 × 7.8 × 7.8 mm^3^ and covered each full cross section of the samples. X-ray source settings were 60 kV and 240 µA. 1000 images were recorded during the 360 ° rotation of the sample. Then, the paraffin was broken at the ends in order to allow the water flow and the sample was set in a Cavitron during 5 min at 0.08 MPa, immerged in paraffin and scanned again with the microtomograph at the same location than previously to observe the new embolism status. The same procedure was repeated for increasing pressure steps, until - 4 MPa (Fig. 1).

**Figure 1:**
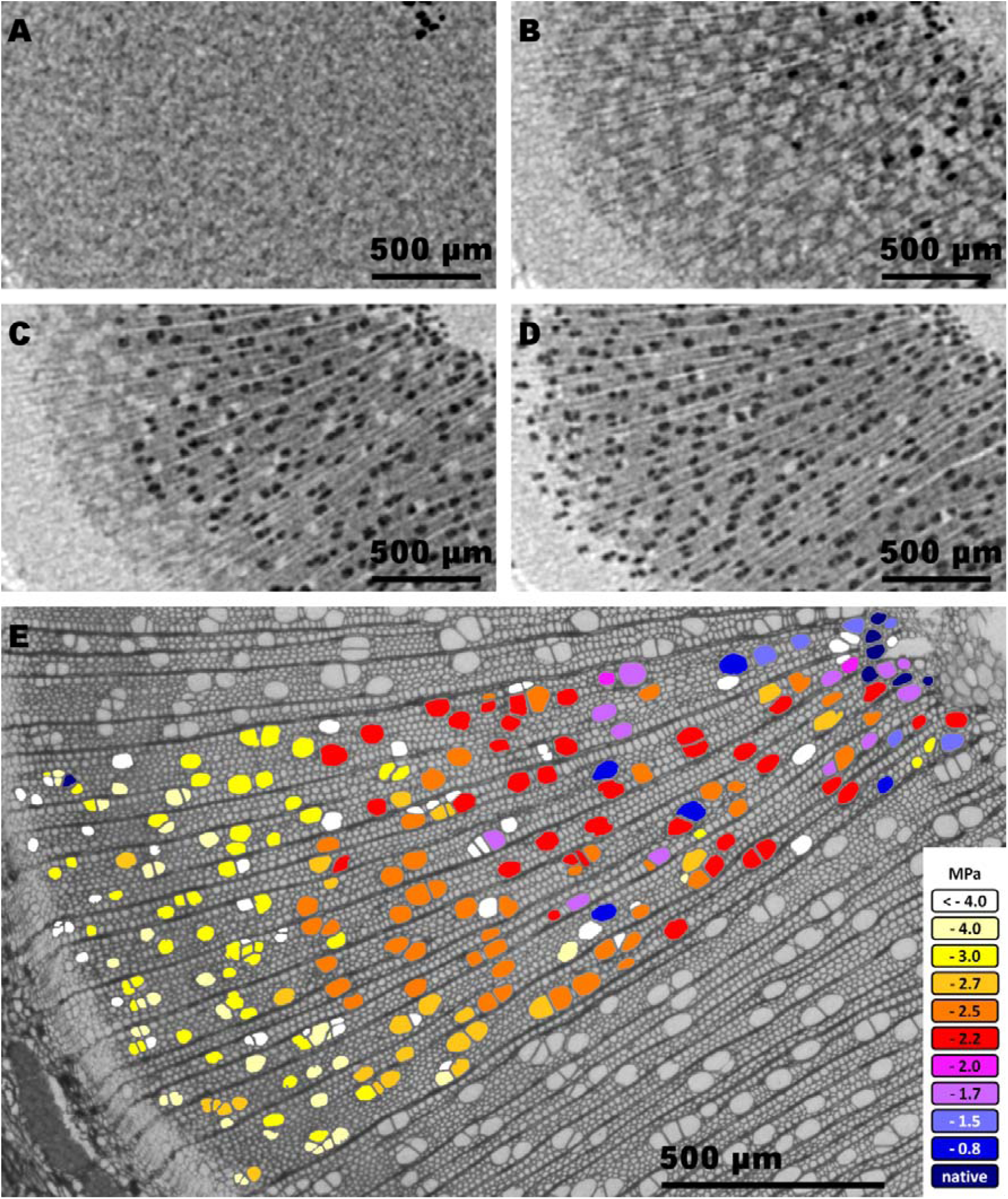
Measurement of the embolism pressure (*P*_e_) of each individual vessel. A-D: Direct observation of embolism spread using a x-ray microtomograph in an intact xylem stem under increasing tension. Black areas reveal the embolized vessels. A: native state (Ψ = 0 MPa). B: Ψ = - 1.5 MPa. C: *P*_50_ state (Ψ = - 2.5 MPa). D: final state (Ψ = - 4 MPa). E: Cut of the same stem sample observed using light microscopy. The resulting image resolution allows us measuring accurately the anatomical traits. Colour represents the embolism pressure (*P*_e_) of each vessel, as measured with x-ray microtomography. Shown images are from a subset of approx. 230 vessels on a Control plant.

Then, the stem sample was cut in the air at 5 mm above the scanned section in order to embolize 100 % of the functional vessels and a last microtomography scan was performed in order to visualize the complete vessel network.

The sample was then dried several days in room conditions and a 25 µm thick cross section slice was cut with a rotary microtome (RM2165, Leica Microsystems). Sections were dyed with series of baths as following: bleach (about 15 sec), acetic acid, Astra blue (1 min), acetic acid, safranin (1 min), acetic acid with a water bath between each solution, then an ethanol series (50, 70, 100 and 100 %). The sections were mounted in Eukitt. Images were processed using a microscope (Zeiss Axio Observer Z1), a digital camera (AxioCam MRc) and Zen imaging software system (Zeiss, Jena, Germany). Image analyses were performed using Fiji software (under ImageJ version 2.0.0-rc-68/1.52h) (Schindelin et al. 2012, Schneider et al. 2012). Diameter of each vessel 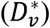 was estimated as the diameter of the circle with the same area as the vessel lumen. For each vessel, the number of vessel in the group (Group Size; GS) and the fraction of membrane in contact with other vessels 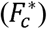 was also measured.

The microtomography scans were reconstructed in three-dimension (3D) using Phoenix datosx 2 software (General Electric, Boston, MA, USA) with spatial resolution of 6.8 **×** 6.8 **×** 6.8 µm^3^ per voxel. Then, for each 3D-reconstruction, a cross section was extracted at the exact same location as with the microscopy section. Using the series of x-ray scans, the embolism pressure in a vessel (*P*_e_) was defined as the pressure step from which the vessel appeared to be air-filled.

Images from microtomography observation (virtual cross sections built by 3D reconstruction) and microscopy observation (stem cross section observed by light microscopy) were aligned using the “Align image by line ROI” tool of Fiji software. A unique identification number was given to each vessel observed in images from both techniques, in order to link the embolism pressure with anatomical parameters (Fig. 1, E). A total of 2570 vessels were identified. Vessels were grouped per 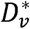, per 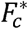 and per GS classes. Classes were sized to be as uniform as possible, counting from 183 up to 748 vessels. A total of 1100 solitary vessels were grouped in the same class when required. Cumulative number of embolized vessels were plotted according to their *P*_e_ and, for each class, a Weibull function was fit (equation 1).

### Statistical analysis

The statistical analysis was performed using the RStudio software (version 1.1.456; running under R core version 3.5.1, R Development Core Team 2008). One way ANOVA were used for comparing the means between the three growth conditions. When we found a significant difference, we referred to Tukey’s multiple range test at P < 0.05 to compare the mean values between growth conditions. The correlation between the structural traits and the *P*_50_ and *P*_e_ were calculated using linear regressions.

## Results

Continuous recordings of the radial growth showed a significant lower growth for the Droughted plants (Table 2). These plants also showed a lower height, lower leaf area, lower *Ψ*_pd_ and lower *Ψ*_mid_, demonstrating that these plants grown under a constrained water regime were affected for their development when compared to Control and Shaded plants. The higher leaf area for Shaded plants compared to Control plants is an evidence that the shading conditions affected the plant development.

**Table 2:** Physiological characterisation of sapling grown under the three different conditions.

| Factor | Unit | Droughted | Control | Shaded |
| --- | --- | --- | --- | --- |
| $\Psi_{pd}$ | Mpa | $-0.59 \pm 0.44^a$ | $-0.14 \pm 0.03^b$ | $-0.11 \pm 0.02^b$ |
| $\Psi_{min}$ | MPa | $-1.44 \pm 0.33^a$ | $-0.98 \pm 0.06^b$ | $-0.98 \pm 0.11^b$ |
| LA | cm<sup>2</sup> | $87.64 \pm 18.41^a$ | $137.61 \pm 17.55^b$ | $184.13 \pm 40.34^c$ |
| Height | mm | $1685 \pm 187^a$ | $2237 \pm 263^b$ | $2282 \pm 76^b$ |
| Diameter | mm | $9.47 \pm 0.83^a$ | $13.96 \pm 0.49^b$ | $10.94 \pm 0.61^b$ |
| $K_h$ | mmol.s<sup>-1</sup>.MPa<sup>-1</sup>.m<sup>-1</sup> | $527.21 \pm 186.63^a$ | $766.92 \pm 154.71^b$ | $780.60 \pm 110.01^b$ |
| $K_s$ | mmol.s<sup>-1</sup>.MPa<sup>-1</sup>.m<sup>-1</sup> | $585.65 \pm 106.76^a$ | $579.50 \pm 167.38^a$ | $529.88 \pm 183.41^a$ |
| $P_{50}$ | MPa | $-3.03 \pm 0.23^a$ | $-2.49 \pm 0.10^b$ | $-2.27 \pm 0.18^b$ |
| $P_{12}$ | MPa | $-2.55 \pm 0.34^a$ | $-2.02 \pm 0.11^b$ | $-1.87 \pm 0.11^b$ |
| $P_{88}$ | MPa | $-3.51 \pm 0.24^a$ | $-2.95 \pm 0.11^b$ | $-2.68 \pm 0.11^b$ |
| Native Embolism | % | $1.81 \pm 10.47^a$ | $-7.48 \pm 7.91^{ab}$ | $-10.81 \pm 7.84^b$ |

Data are mean values ± standard deviation for each growth condition. For each line, values not followed by the same letter differ significantly at $p < 0.05$ (one-way ANOVA). $\Psi_{pd}$ , Predawn water potential; $\Psi_{min}$ , Minimum water potential; LA, Leaf area; $K_h$ , Theoretical hydraulic conductivity; $K_s$ , Specific hydraulic conductivity; $P_{50}$ ; $P_{12}$ ; $P_{88}$ , pressure inducing 50; 12 and 88 percent loss of conductance.

Growing clone plants under different environmental conditions aimed to induce wide variations in xylem vulnerability to embolism. The three growth conditions spread the measured *P*_50_ in a range from - 2.00 to - 3.47 MPa. A significantly lower *P*_50_ was found on Droughted when compared to Control and Shaded plants (*p-value < 0.001*, Table 2), while the slopes of the vulnerability curves were not different depending on the growth conditions (Fig. 2, A). Despite a slightly higher native embolism measured on Droughted plants compared to Shaded plants, *Ψ*_mid_ was higher than the inflexion point of the vulnerability curve (*P*_12_) for each growth condition. This allows excluding any effect of these quite low native embolism on measured *P*_50_. There was no difference on mean *K*_S_ between the growth conditions (Fig. 2, B), suggesting no plasticity for this trait in our experimental conditions. When considering the vessel diameter measured by light microscopy, a reduced *K*_h_ was measured in the Droughted plants compared to Control and Shaded plants (Table 2).

**Figure 2:**
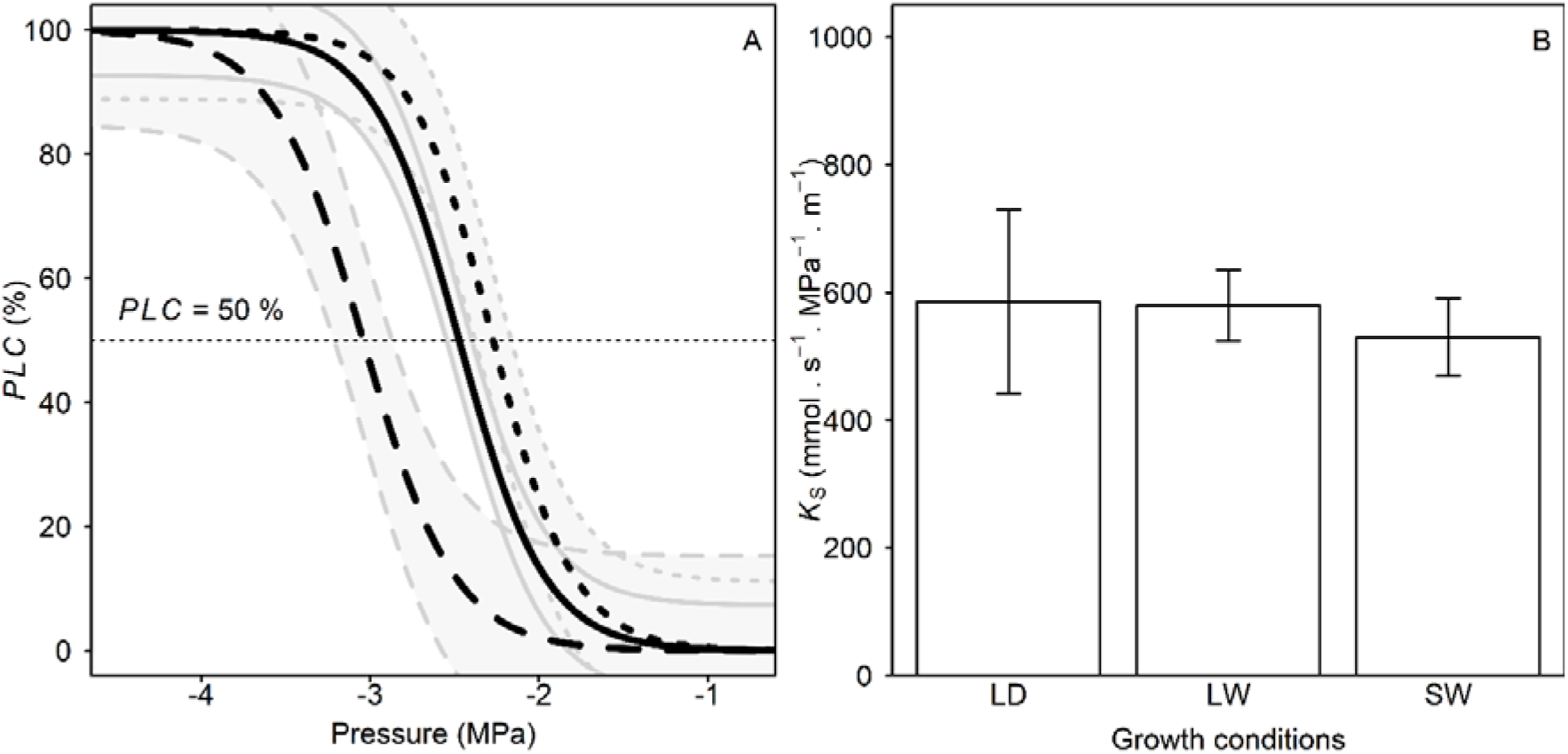
Xylem hydraulic traits in trees depending on the growth conditions. A: Xylem vulnerability to embolism curve. Each line is the mean curve per condition: Droughted, n = 9 from 9 trees; Control, n = 10 from 5 trees; Shaded, n = 12 from 6 trees. Dashed line, Droughted plants; full line, Control plants; dotted line, Shaded plants. Grey areas represent the standard deviations around the means. Horizontal dotted line indicates the 50 % loss of conductance. B: Hydraulic specific conductivity (*K*_s_). Data are mean values for 8 Droughted trees, 9 Control trees, 9 Shaded trees. Error bars show the standard deviation.

The analyses combining diverse observation methods (light microscopy, TEM, SEM), allowed measuring a large set of anatomical traits from tissue to pit level. The correlation between these traits and the *P*_50_ was assessed (Fig. 3, 4).

**Figure 3:**
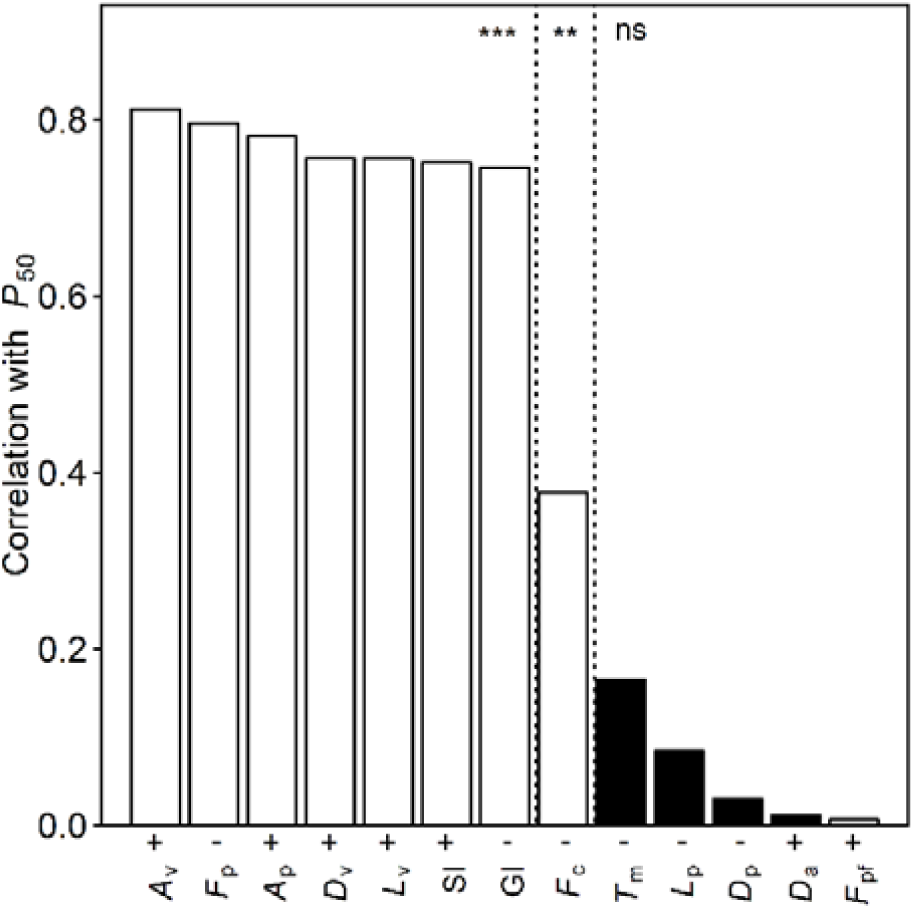
Correlation between *P*_50_ and several xylem structural traits. Data are squares of the coefficient of correlation (*R*^*2*^) for each factor with *P*_50_. Black bars indicate pit-related traits and white bars indicate vessel and xylem-related traits. On the x-axis, a “+” symbol indicates a positive correlation, while a “-” symbol indicates a negative one. Stars indicate the significance of the correlation: “***”, *p-value* < 0.001; “**”, *0.001 < p-value < 0.01*; “ns”, non-significant correlation.

**Figure 4:**
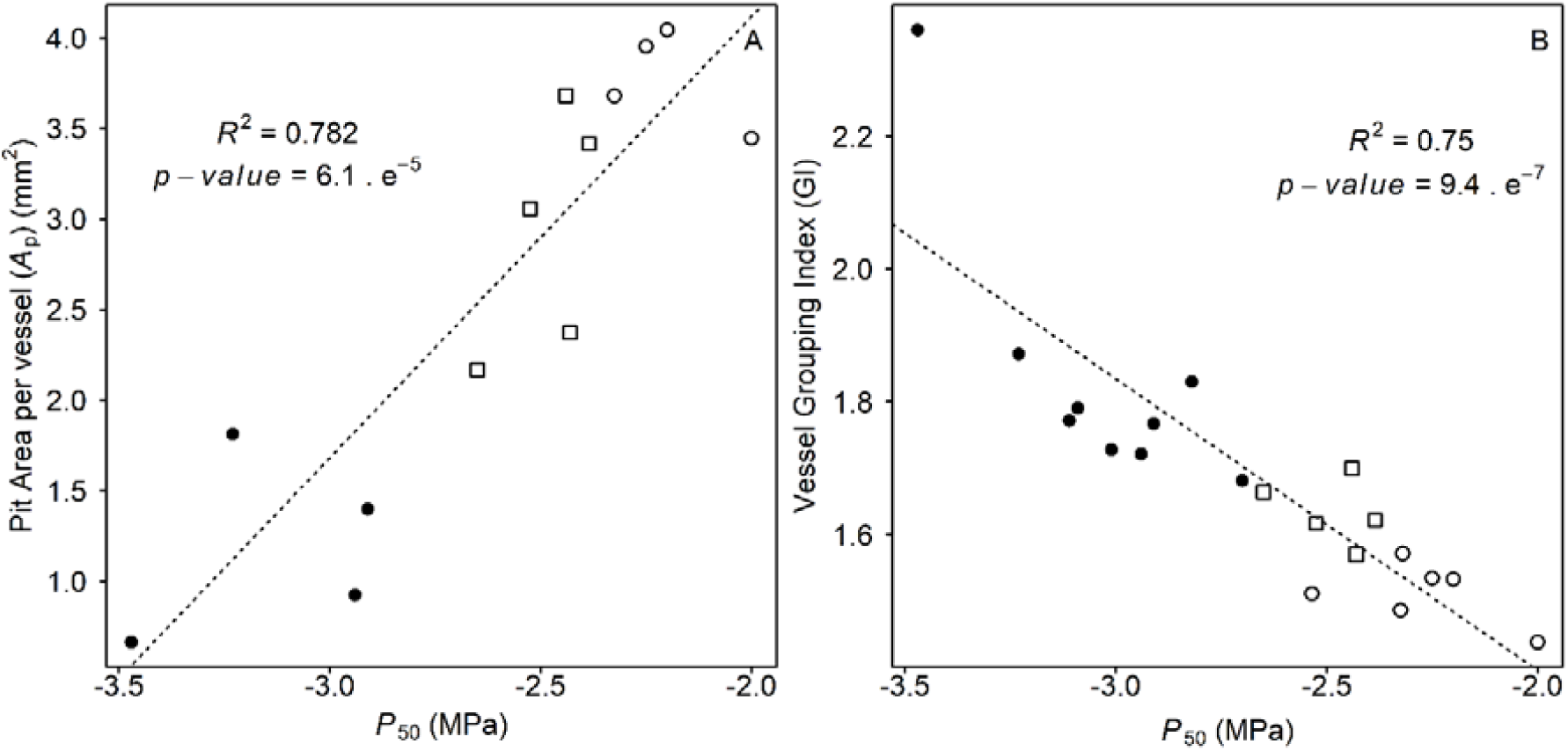
Correlation between *P*_50_ and two xylem structural traits. A: Relationship between *P*_50_ and pit area per vessel (*A*_p_). B: Relationship between *P*_50_ and vessel grouping index (GI). Each point represents the mean value for an individual tree. Black circles refer to Droughted plants; white circles refer to Control plants and white squares refer to Shaded plants. The dotted line is the regression line.

The traits measured at tissue level (GI, SI and *F*_p_) showed a strong linear correlation with *P*_50_ (*R*^*2*^ *> 0.70; p-value < 0.001*; Fig. 3, 4), except *F*_c_ that exhibited a weaker correlation (*R*^*2*^ = 0.377; *p-value* = 0.0040). These results put in light a relationship between vessels connectivity and grouping and vulnerability to embolism (negative relationship for *F*_c_, GI and *F*_p_; positive relationship for SI). However, we also found no correlation between Pit-field fraction (*F*_pf_) and *P*_50_, with no variation among the growth conditions (Table 3). We observe a strong positive relationship (*p-value < 0.001*) between *P*_50_ and the vessel dimensions (*L*_v_, *D*_v_ and *A*_v_) showing that larger vessels with larger pit area tend to be associated with an increase in vulnerability to embolism (*R*^*2*^ *> 0.75; p-value < 0.001*). The positive correlation between *P*_50_ and *A*_p_ (*R*^*2*^ *= 0.782; p-value < 0.001*, Fig. 4) enlightened the link between the area of vessel covered by bordered pits and the xylem vulnerability to embolism.

**Table 3:** Xylem structural traits depending on the growth conditions.

| **Trait** | **Unit** | **Droughted** | **Control** | **Shaded** |
| --- | --- | --- | --- | --- |
| $A_p$ | mm<sup>2</sup> | 1.20 ± 0.51<sup>a</sup> | 2.94 ± 0.65<sup>b</sup> | 3.78 ± 0.27<sup>c</sup> |
| $A_v$ | mm<sup>2</sup> | 8.21 ± 3.93<sup>a</sup> | 20.63 ± 3.64<sup>b</sup> | 26.65 ± 8.22<sup>b</sup> |
| $D_a$ | μm | 3.67 ± 0.34 | 3.37 ± 0.61 | 3.98 ± 0.81 |
| $D_p$ | μm | 9.18 ± 0.69 | 8.64 ± 0.55 | 8.89 ± 0.72 |
| $D_v$ | μm | 31.22 ± 6.14<sup>a</sup> | 40.07 ± 1.98<sup>b</sup> | 42.71 ± 2.28<sup>b</sup> |
| $F_c$ | % | 20.35 ± 2.70<sup>a</sup> | 19.01 ± 1.13<sup>b</sup> | 17.04 ± 0.96<sup>c</sup> |
| $F_p$ | % | 15.95 ± 1.13<sup>a</sup> | 14.15 ± 0.77<sup>b</sup> | 12.53 ± 0.56<sup>c</sup> |
| $F_{pf}$ | % | 74.45 ± 1.33 | 74.46 ± 2.93 | 74.13 ± 2.75 |
| **GI** | - | 1.84 ± 0.20<sup>a</sup> | 1.63 ± 0.05<sup>b</sup> | 1.51 ± 0.05<sup>c</sup> |
| $L_p$ | μm | 1.99 ± 0.06 | 2.08 ± 0.03 | 1.87 ± 0.11 |
| $L_v$ | mm | 70.79 ± 25.1<sup>a</sup> | 137.0 ± 18.86<sup>b</sup> | 164.6 ± 48.4<sup>c</sup> |
| **SI** | % | 33.13 ± 5.43<sup>a</sup> | 38.46 ± 2.19<sup>b</sup> | 43.73 ± 3.11<sup>c</sup> |
| $T_m$ | μm | 0.26 ± 0.04 | 0.23 ± 0.04 | 0.24 ± 0.02 |

No linear correlation appeared between the qualitative pit parameters (*D*_a_, *D*_p_, *L*_p_ and *T*_m_) and the *P*_50_: we observed no variation for *D*_a_, *D*_p_ and *T*_m_ among growth conditions.

The direct microtomographic visualization of embolism inside the xylem (Fig. 1) allowed evaluating the vulnerability to embolism of individual vessels classified depending on their structural parameters (Fig. 5). The correlation between 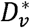 and *P*_e_ (Fig. 5, A) was clear: wider vessels appeared more vulnerable than the narrower ones. 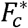 showed a smaller influence on *P*_e_ (Fig. 5, B): solitary vessels 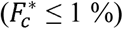 and weakly connected vessels 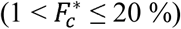 were more vulnerable than the highly connected vessels 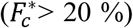. The link between GS and *P*_e_ (Fig. 5, C) appeared to be the less clear: the most vulnerable vessels were the solitary ones whereas the grouped vessels (GS ≥ 2) were less vulnerable. Despite a significant correlation between *P*_e_ and 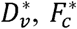 and GS (*p-value < 0.001;* Fig. 5, D), the strength of the correlation was very poor (*R*^2^ < 0.25).

**Figure 5:**
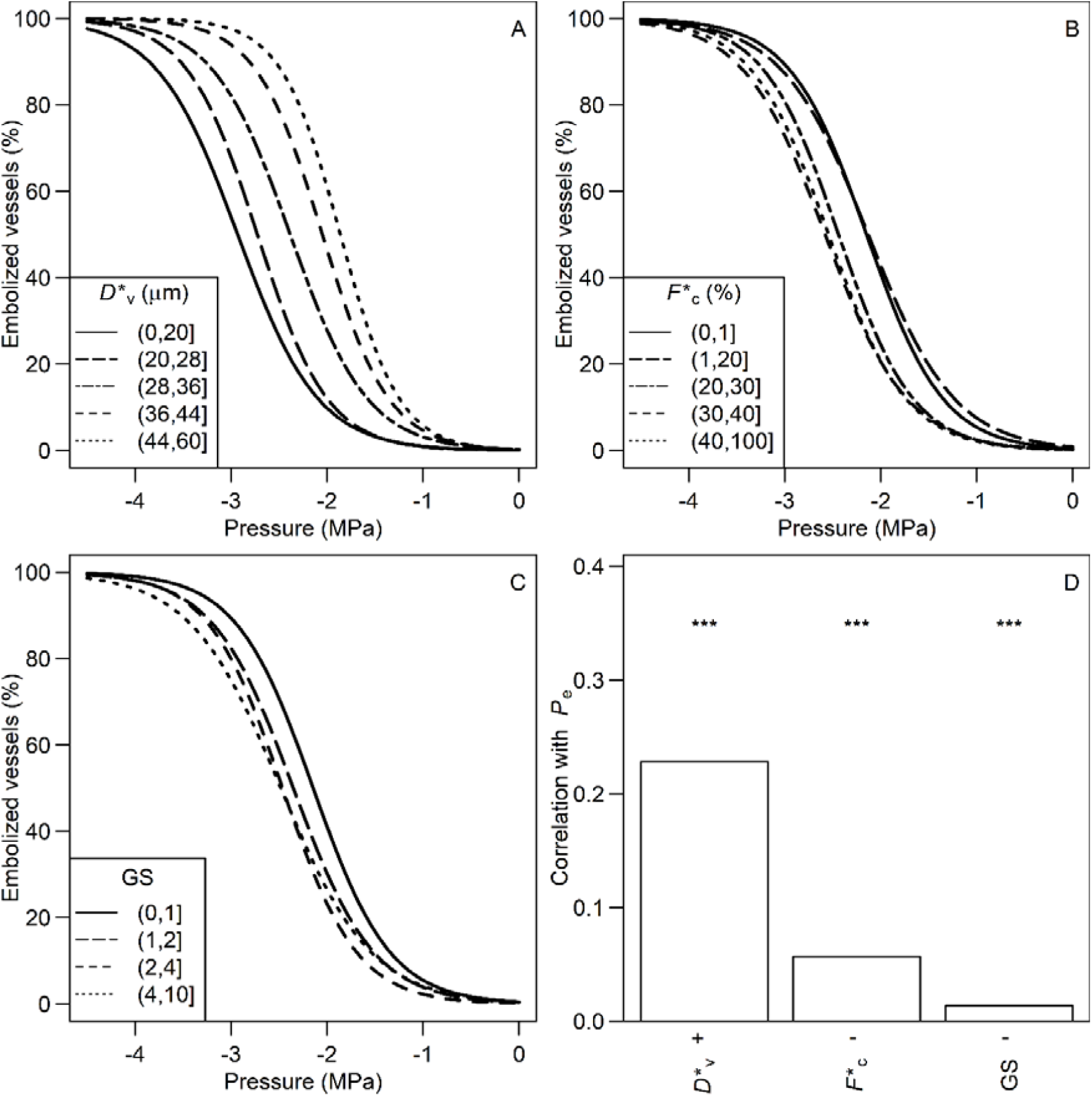
Correlation between *P*_e_ and vessel traits inside a xylem. Data are all vessel measurements pooled from analyses on four individuals using X-ray microtomography. A-C: Vulnerability to embolism curves of vessels grouped by classes depending on structural traits. A: Vessels clustered by diameter 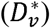 classes. The dash sizes of the lines indicate the vessel diameter class: from full line (narrow vessels) to dotted line (wide vessels). B: Vessels clustered by classes for fraction of membrane length in contact with other vessels 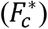. The dash sizes of the lines indicate the vessel contact fraction class: from full line (non-contact vessels) to dotted line (vessels sharing high portion of membrane length). C: Vessels are clustered by group size (GS) classes. The dash sizes of the lines indicate the vessel group sizes: from full line (solitary vessels) to dotted line (vessels in large groups). D: Correlation between *P*_e_ and xylem structural traits. Data are squares of the coefficient of correlation (*R*^*2*^) for each factor with *P*_e_. On the x-axis, a “+” symbol indicates a positive correlation, while a “-“ symbol indicates a negative one. Stars indicate the significance of the correlation for the trait: “***”, *p-value* < 0.001.

## Discussion

The range for *P*_50_ plasticity induced by the growth conditions was large: 0.76 MPa between the mean *P*_50_ of Droughted and Shaded plants (Table 2; Fig. 2, A) and up to 1.47 MPa between two individuals. This is consistent with previous studies: Awad et al. (2010) got a difference of 0.63 MPa between droughted and well-watered plants; Plavcová and Hacke (2012) go a difference of 1.08 MPa between droughted and shaded *Populus trichocarpa x deltoides* plants. In this latter study, they reported a variation of *P*_50_ up to 1.56 MPa but their setup also included treatment in nutrient availability, and they recorded r-shaped curves that cannot be compared with our S-shaped curves. Therefore, the plasticity induced by our experimental setup was probably close to the maximum we could expect according to the literature.

The absence of difference in specific hydraulic conductivity (*K*_s_) between, Droughted and Control plants (Table 2; Fig. 2, B) not surprising since poor correlation between vulnerability to embolism and *K*_s_ has been reported in a meta-analysis (Gleason et al. 2016). Furthermore, the lack of trade-off between hydraulic efficiency and safety was observed within species (Awad et al. 2010, Plavcová and Hacke 2012, Schuldt et al. 2016). A significant decrease of the theoretical conductivities (*K*_h_) was found for Droughted plants compared to other plants (Table 2), relying on a decrease in vessel diameter (*D*_v_) (Table 3); whereas the pit structure was not modified (Table 3). The decrease in lumen conductance in Droughted plants may be offset by other changes we did not investigate, such as vessel wall carving, pit biochemistry or pit membrane porosity.

*P*_50_ was correlated with anatomical traits related to quantitative pit characteristics measured at the xylem and vessel levels (significant correlations with *R*^2^ > 0.7 for 7 out of the 9 traits; Fig. 3, 4). By contrast, no correlation was found with the traits related to the qualitative pit characteristics we measured, i.e. the pit dimensions (*D*_a_, *D*_p_, *L*_p_ and *T*_m_; Fig. 3). Thus, the pit ultrastructure does not appear as a driver of the plasticity of vulnerability to embolism in *Populus tremula x alba*. The pit membrane pore sizes contribute to the differences in vulnerability to embolism (Jansen et al. 2009); but this parameter is difficult to measure accurately because pores include a series of various pore constrictions, and the most narrow constriction will be the main bottleneck. The role of the biochemical composition of the pit membrane in vulnerability to embolism plasticity cannot be excluded. Once again, pit biochemistry was investigated using immunolabelling (Kim et al. 2011, Herbette et al. 2015), but accurate techniques are needed to investigate within-species difference in *P*_50_. Moreover, calcium in pit membrane was reported to be a major determinant of between-species differences in vulnerability to embolism, but it was not involved in the plasticity of vulnerability to embolism (Herbette and Cochard 2010).

Pit ultrastructure, especially the pit membrane thickness were identified as the major traits involved in variation in vulnerability to embolism between species (Jansen et al. 2009; Tixier et al. 2014). In addition, between species differences in vulnerability to embolism also depend on pit mechanical behaviour (Tixier et al. 2014). The probability for air seeding through large pores is expected to be higher when more pits are present (rare pit hypothesis proposed by Christman et al. 2009). The pit area can thus explain differences in vulnerability to embolism among some angiosperm groups but not others (Lens et al. 2013). This trait, which depends on the vessel dimensions and xylem organization, does not appear very relevant to explain variability in vulnerability to embolism between species. Lens et al. (2011) tested the relationship between several qualitative and quantitative pit properties and vulnerability to embolism for 11 acer species. They found that vulnerability to embolism strongly correlated with depth of bordered pit chamber (*L*_p_) and pit membrane thickness (*T*_m_) whereas no relationship was found between vulnerability to embolism and vessel diameter (*D*_v_) and total pit area per vessel (*A*_p_). By contrast, our results suggest that the plasticity of vulnerability to embolism plasticity is controlled by the xylem organization and vessel dimensions, and not by changes in pit structure. Thus, the mechanisms controlling the inter-specific variability in vulnerability to embolism seem to be different from the drivers of the within species plasticity.

Vulnerability curves are commonly established by measuring the impact of embolism on the conductance, but not by measuring the embolism rates. Because an embolized vessel can induce different effects on the xylem conductance and thus the value of *P*_50_, depending on vessel size and xylem organisation, “hydraulic vulnerability” would be a more suitable term when we compare xylems for *P*_50_ using methods based on hydraulic measurements. That is why we will use now this term in the following lines. X-ray microtomography investigations allow the visualisation of embolized vessels, and not the loss of hydraulic conductance. Thus, it really allows measuring the vulnerability to embolism and not the hydraulic vulnerability.

Our results showing a strong relationship between *P*_50_ and some vessel and xylem parameters provide three non-exclusive explanations for the acclimation of hydraulic vulnerability. This latter relies on changes in vulnerability to embolism of the vessels or on changes in the effect of the embolism on conductance. First, our study shows that vulnerable individuals exhibited bigger vessels (both longer (*L*_v_) and wider (*D*_v_); Fig. 3). When a large vessel embolizes, it generates a greater impact on the hydraulic conductivity compared to a smaller vessel. Thus, a xylem having a high proportion of large vessels undergoes an important drop of conductivity after each vessel embolism. Second, we found that vulnerable xylems had a greater SI and a lower GI and *F*_c_. Redundancy in the xylem has already been linked with a lower hydraulic vulnerability using modelling (Ewers et al. 2007). High connectivity and grouping is an efficient way to maintain the hydraulic conductance despite embolized vessels in the xylem by providing alternative pathways to the water flow (Carlquist 1966, Schuldt et al. 2016). Third, larger vessels have a larger pit area per vessel (*A*_p_) and would thus be more prone to embolism, according to the pit area hypothesis (Christman et al. 2009). Direct observations using X-ray microtomography allowed monitoring the dynamics of xylem embolism and in particular to determine the embolism pressure for each vessel in a stem sample (Fig.1; Fig. 5). This approach supports the third explanation, since larger vessels 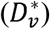 had a higher vulnerability to embolism– as noticed by Cai and Tyree (2010) using a statistical and indirect and destructive technique. Nevertheless, the poor correlation (low *R*^2^ values) between the embolism pressure of each vessel (*P*_e_) and 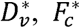 or GS give us clue that the rare pit hypothesis is far from being sufficient for explaining the hydraulic vulnerability inside a stem sample. Other additional mechanisms would be involved to explain the plasticity of hydraulic vulnerability observed among growth conditions: they would include the effect of redundancy and of vessel embolized volume on the loss of conductance. That is why we assume that the different mechanisms we described here act together to design the hydraulic vulnerability during acclimation.

In conclusion, we found that the acclimation of vulnerability to embolism to contrasted growth conditions occurs without any change in pit ultrastructure, contrary to what was reported when comparing species. Thus, within-species plasticity and between-species variability for vulnerability to embolism rely on different mechanisms. Instead, we showed that an increase in resistance to embolism in poplar is related to an increase in vessels connectivity and grouping and a decrease in vessel dimensions, leading to reduce the likely of air seeding through a pit in a vessel and the effect of such embolism events on hydraulic conductance. This study will allow focusing on the relevant candidate genes controlling vulnerability to embolism such as those involved in vessels grouping and connectivity or vessel dimensions. These genes include the aquaporins involved in cell expansion during xylogenesis (Plavcová et al. 2013), the genes controlling the cell wall metabolism in xylem such as *VND6, VND7* and *MYB46*, which expression levels changed in response to an abiotic stress (Plavcová et al. 2013, Taylor-Teeples et al. 2016) or *CLE* genes (*CLE41* and *CLE44*) that repress the xylem differentiation (De Rybel et al. 2016).

## Acknowledgements

The authors thanks Christelle Boisselet and Brigitte Girard for the plant production, Christophe Serre for the LDVT preparation, Romain Souchal for the balance and LVDT installation, Patrice Chaleil, Aline Faure and Stephane Ploquin for watching the plants in the greenhouse, André Marquier for the PAR measurements and Felix Hartmann for his help in calculating the vessel length.

## Authors’ contributions

C.L. and S.H. designed the study and wrote the manuscript with contributions from all authors. C.L., P.C. and J.C. performed field work and hydraulic measurements; C.L., N.B-M., Y.Q., L.B. and J.S. performed electron microscopy; C.L., N.B-M., P.C. performed light microscopy; C.L., P.C., E.B. and J.C. performed X-ray microCT; C.L., P.C. and E.B. performed image analysis. All authors approved this manuscript.

